# Ultra-fast and accurate motif finding in large ChIP-seq datasets reveals transcription factor binding patterns

**DOI:** 10.1101/394007

**Authors:** Yang Li, Pengyu Ni, Shaoqiang Zhang, Guojun Li, Zhengchang Su

## Abstract

The availability of a large volume of chromatin immunoprecipitation followed by sequencing (ChIP-seq) datasets for various transcription factors (TF) has provided an unprecedented opportunity to identify all functional TF binding motifs clustered in the enhancers in genomes. However, the progress has been largely hindered by the lack of a highly efficient and accurate tool that is fast enough to find not only the target motifs, but also cooperative motifs contained in very large ChIP-seq datasets with a binding peak length of typical enhancers (∼ 1,000 bp). To circumvent this hurdle, we herein present an ultra-fast and highly accurate motif-finding algorithm, ProSampler, with automatic motif length detection. ProSampler first identifies significant *k*-mers in the dataset and combines highly similar significant *k*-mers to form preliminary motifs. ProSampler then merges preliminary motifs with subtle similarity using a novel graph-based Gibbs sampler to find core motifs. Finally, ProSampler extends the core motifs by applying a two-proportion *z*-test to the flanking positions to identify motifs longer than *k*. As the number of preliminary motifs is much smaller than that of *k*-mers in a dataset, we greatly reduce the search space of the Gibbs sampler compared with conventional ones. By storing flanking sequences in a hash table, we avoid extensive IO and the necessity of examining all lengths of motifs in an interval. When evaluated on both synthetic and real ChIP-seq datasets, ProSampler runs orders of magnitude faster than the fastest existing tools while more accurately discovering primary motifs as well as cooperative motifs than do the best existing tools. Using ProSampler, we revealed previously unknown complex motif occurrence patterns in large ChIP-seq datasets, thereby providing insights into the mechanisms of cooperative TF binding for gene transcriptional regulation. Therefore, by allowing fast and accurate mining of the entire ChIP-seq datasets, ProSampler can greatly facilitate the efforts to identify the entire *cis*-regulatory code in genomes.

## INTRODUCTION

Although different types of somatic cells in a multicellular organism contain the same genome, they produce a unique set of gene products for their specific functions. This is achieved by gene transcriptional regulation during the course of embryogenesis and development, as well as in response to the environmental changes. Gene transcriptional regulation is mainly carried out by interactions between transcription factors (TF) and specific DNA sequences called TF binding sites (TFBSs).TFBSs are typically short DNA segments with a length of 6-20 base pairs (bp). TFBSs recognized by the same TF are highly similar and are collectively called a *motif*. Identifying all TFBSs of all TFs in a genome (called the regulatory lexicon(Deplancke et al. 2016)) is a central but highly challenging task in this post-genomic era. Fortunately, the rapid development of next-generation-sequencing (NGS) based high-throughput technologies has largely facilitated the efforts to identify all the TFBSs in sequenced genomes. Particularly, chromatin immunoprecipitation (ChIP) followed by sequencing (ChIP-seq)(Park 2009) and its derivative (e.g. MNChIP-seq(Tsankov et al. 2015)) have become the mainstream methods for genome-scale characterization of TFBSs for target TFs.

In a ChIP-seq experiment, one can obtain tens or even hundreds of thousands of binding peaks of the ChIP-ed/target TF in a tissue/cell sample, with a length from hundreds to thousands bp returned by a peak-calling tool based on the sequenced reads mapped to the reference genome(Valouev et al. 2008; Zhang et al. 2008). Although TFBSs of the target TF are usually enriched, identification of all the TFBSs in such a very large number of binding peaks is still a highly challenging task. First, the sheer large volume of sequences in a dataset dwarfs existing classic motif-finding tools(Bailey and Elkan 1994; Pavesi et al. 2001; Fauteux et al. 2008) that are mainly aimed at datasets of a small size derived from co-expressed genes(Prakash and Tompa 2005), or phylogenetic foot-printing analyses(Blanchette et al. 2002; Pavesi et al. 2007). Consequently, in many ChIP-seq studies, only a few hundred of top-scored binding peaks were used for motif-finding(Machanick and Bailey 2011; Kheradpour and Kellis 2013), which clearly under-exploited the valuable datasets. Although new faster algorithms such as Trawler(Ettwiller et al. 2007), ChIPMunk(Kulakovskiy et al. 2010), HMS(Hu et al. 2010), CMF(Mason et al. 2010), Homer(S et al. 2010), STEME(Reid and Wernisch 2011), DREME(Bailey 2011), DECOD(Huggins et al. 2011), RSAT(Thomas-Chollier et al. 2012), POSMO(Ma et al. 2012), XXmotif(Hartmann et al. 2013), Dimont(Grau et al. 2013), and FastMotif(Hartmann et al. 2013), motifRG(Yao et al. 2014), EXTREME(Quang and Xie 2014) and WSMD(Zhang et al. 2017) have been developed using various algorithmic approaches, most of these tools are still too slow, thus cannot work on very large ChIP-seq datasets from mammalian tissues.Second, most of these tools are aimed to identify the motif of the target TF in shortened binding peaks for higher computational efficiency and accuracy(Huggins et al. 2011; Ma et al. 2012; Thomas-Chollier et al. 2012; Grau et al. 2013; Hartmann et al. 2013; Jia et al. 2013; Colombo and Vlassis 2015; Ikebata and Yoshida 2015) and some even depends on the TF binding information(Ma et al. 2012; Grau et al. 2013). Nonetheless, since TFBSs of cooperative TFs tend to be closely located, forming *cis*-regulatory modules (CRMs) such as enhancers and promoters, there is increasing interest in identifying motifs of cooperative TFs in addition to the motif of the target TF (*primary motif*) in longer or extended binding peaks with a length close to those of typical enhancers (∼ 1,000 bp)(Gerstein et al. 2010; Chen and Zhou 2011; Negre et al. 2011; Whitington et al. 2011; Zhang et al. 2011; Bailey and Machanick 2012; Sun et al. 2012). Third, faster tools (e.g., Homer, DECOD, RSAT and DREME(Bailey 2011), POSMO) are based on the discriminative motif-finding schema(Sinha 2003) by finding overrepresented *k*-mers in a ChIP-seq dataset, but they often fail to identify TFBSs with subtle degeneracy(S et al. 2010; Bailey 2011), thus there are still large rooms for improvement in sensitivity and specificity. Fourth, most existing motif-finding tools return too many false positive motifs, making it difficult for the user to decide which ones are likely to be authentic, whereas some other existing tools are even unable to determine the number of motifs in a dataset, requiring the user to specify the number of motifs to be found, which is unknown in almost all applications. Finally, most current motif finding algorithms can only identify motifs with a pre-specified length, while those that are able to determine the length of motifs employ an exhaustive enumeration strategy within an interval of length, requiring large memory and running time(Pavesi et al. 2001; S et al. 2010; Bailey 2011; Machanick and Bailey 2011).

In order to circumvent these obstacles, we have developed an ultrafast and accurate motif-finding algorithm and tool named ProSampler (<u>Pro</u>file <u>Sampler</u>) with the ability to automatically determine the number of motifs in a dataset and the length of each motif, using a combination of discriminative heuristic seeding(Bailey 2011; Hartmann et al. 2013), Gibbs sampling(Lawrence et al. 1993; Liu et al. 2001) and length extension. More specifically, ProSampler first numerates all nontrivial *k*-mers in the binding peaks while tracking the *k*-mers’ flanking *l*-mers. It then combines highly similar nontrivial *k*-mers, forming a set of preliminary motifs. ProSampler then identifies *k*-mer core motifs by merging similar preliminary motifs using a Gibbs sampler on a graph that reflects the similarity of the preliminary motifs. Finally, ProSampler finds the final motifs by extending the core motifs to the flanking regions using a statistical test. When evaluated on both synthetic and real MNChIP-seq datasets with various binding peak lengths (200, 500 and 1,000 bp), ProSampler is orders of magnitude faster than six existing fastest motif-finding tools, including BioProspector(Liu et al. 2001), DREME(Bailey 2011), XXmotif(Hartmann et al. 2013), Homer(S et al. 2010), motifRG(Yao et al.2014) and Dimont(Grau et al. 2013), while identifying all the implanted motifs in the synthetic datasets with the highest specificity, and more primary motifs as well as cooperative motifs in the real datasets. The source code, executives and relevant data are available at Github: https://github.com/zhengchangsulab/prosampler.

## RESULTS

### The ProSampler algorithm

First, we numerate all *k*-mers (*k* = 8 by default) in the dataset and the corresponding background dataset while tracking each *k*-mers’ two flanking *l*-mers (*l* = 6 by default), and identify a set of significant *k*-mers with a *z*-score >*α* (*α* = 8 by default) and a set of sub-significant *k*-mers with *z*-score >*β* (*β* = 4.5 by default) using a two-proportion *z*-test. Second, for each significant *k*-mer, we combine it with all sub-significant *k*-mers that differ from the *k*-mer by one nucleotide to form a *preliminary motif*. Third, we construct a graph using the preliminary motifs as nodes, and connecting two preliminary motifs by an edge if their Sandelin-Wasserman (SW) similarity(Sandelin and Wasserman 2004b; Gupta et al. 2007a) is greater than a preset value *γ*(*γ* = 1.80 by default). Fourth, we perform Gibbs sampling on the graph using the preliminary motif with the highest *z*-score as the seed, thereby merging preliminary motifs with subtle similarity into a large one called a *k*-mer *core motif*. We remove the merged preliminary motifs from the graph, and then repeat Gibbs sampling on the remaining graph until all nodes are removed, or a specified number of core motifs are identified (option). Fifth, for each core motif, we extend it by considering the nearest columns that are formed by the two flanking *l*-mers based on a two-proportion *z*-test. Finally, we output the final motifs based on their quality scores after removing the redundant motifs. The details of the algorithm are described in Methods.

### Comparison of the programs on synthetic datasets

We first compared ProSampler with five state-of-the-art motif-finding tools, i.e., BioProspector(Liu et al. 2001), Homer(S et al. 2010), DREME(Bailey 2011), XXmotif(Hartmann et al. 2013) and motifRG(Yao et al. 2014) for their speed and ability to find at least a subset of the binding sites of ten implanted JASPAR motifs (Table S1) with different lengths (8-15 bp) in six synthesized datasets, i.e., D_1_∼ D_6_, containing various numbers of sequences (D_1_: 500, D_2_: 1,000, D_3_: 2,000, D_4_: 5,000, D_5_: 10,000 or D_6_: 20,000) with a length of 1,000 bp. Dimont(Grau et al. 2013) was not included in this evaluation as it requires a ChIP-seq quality score of the binding peaks. The 10 motifs were implanted in each dataset at different occurrence frequencies ranging from 0.1 to1.0 site/sequence to mimic a broad spectrum of cooperation between the ChIP-ed TF and its cooperators (Online Methods). As BioProspector needs the user to specify the number of motifs to be found in a dataset, we let it output 150 motifs in each dataset. For the five other programs that are able to automatically determine the number of motifs to be output in a dataset, we let them output all the motifs they found. As expected, the running time of ProSampler scaled linearly to the size of the datasets, and it was orders of magnitude faster than the second fastest program Homer (Fig. 1A). Remarkably, ProSampler identified all the 10 implanted motifs in datasets D_1_∼ D_5_ by its top 10 motifs (Table 1 and Fig. 1B). In D_6_, ProSampler recovered all the 10 implanted motifs by its top 14. Although XXmotif found all the 10 implanted motifs in each of the six datasets by its top 10 motifs (Table 1 and Fig. 1B), one of the predicted motifs in D_1_ matched two implanted ones. When up to 53 top-ranked motifs are considered, DREME also was able to identify all the 60 implanted motifs, while the three other programs failed to identify all the 60 implanted motifs even when all the returned motifs were considered (Table 1 and Fig. 1B), missing motifs with either low or high occurrence frequencies (Fig. 1C). Hence, ProSampler was the only program able to identify all the implanted motifs occurring at either low or high with high ranks without overlapping. Moreover, ProSampler recovered all the 60 implanted motifs implanted the in the six datasets with the smallest number (65) of predicted motifs (Table 1), achieving the lowest false discovery rate (FDR) of 0.08 (Table 1), substantially outperforming the five other programs in this regarding. The 60 motifs predicted by ProSampler matched more significantly (p<0.01, Wilcoxon rank sum test for-log (*q-value*)) the implanted ones by TOMTOM than those predicted by the all the other five programs (Fig. 1D, see Table S1 for the motif logos).

**Table 1.** Prediction of motifs by the six programs in the synthetic datasets

| Datasets | D<sub>1</sub> (500<sup>a</sup>) |  | D<sub>2</sub> (1,000) |  | D<sub>3</sub> (2,000) |  | D<sub>4</sub> (5,000) |  | D<sub>5</sub> (10,000) |  | D<sub>6</sub> (20,000) |  | Total true motifs found | Total false motifs found | SN | SP | FDR |
| --- | --- | --- | --- | --- | --- | --- | --- | --- | --- | --- | --- | --- | --- | --- | --- | --- | --- |
|  | True motifs<sup>b</sup> | Discovered motifs | True motifs | Discovered motifs | True motifs | Discovered motifs | True motifs | Discovered motifs | True motifs | Discovered motifs | True motifs | Discovered motifs |  |  |  |  |  |
| ProSAMPLER | 10 | 10 | 10 | 10 | 10 | 10 | 10 | 10 | 10 | 10 | 10 | 15 | 60 | 5 | 1.00 | 0.92 | 0.08 |
| BioProspector | 10 | 150 | 10 | 150 | 9 | 150 | 9 | 150 | 8 | 150 | 10 | 150 | 56 | 844 | 0.93 | 0.06 | 0.94 |
| DREME | 10 | 14 | 10 | 20 | 10 | 23 | 10 | 36 | 10 | 48 | 10 | 58 | 60 | 139 | 1.00 | 0.30 | 0.70 |
| XXmotif | 10 | 17 | 10 | 10 | 10 | 11 | 10 | 11 | 10 | 12 | 10 | 11 | 60 | 12 | 1.00 | 0.83 | 0.17 |
| Homer | 9 | 300 | 10 | 300 | 10 | 300 | 10 | 300 | 10 | 300 | 10 | 300 | 59 | 1,741 | 0.98 | 0.03 | 0.97 |
| motifRG | 10 | 40 | 8 | 95 | 10 | 91 | 8 | 133 | 7 | 196 | 9 | 150 | 52 | 653 | 0.87 | 0.07 | 0.93 |

<sup>a</sup> Number of sequences in the dataset. <sup>b</sup> Numbers in bold type represent the best performance and efficiency in the column.

**Fig. 1.**
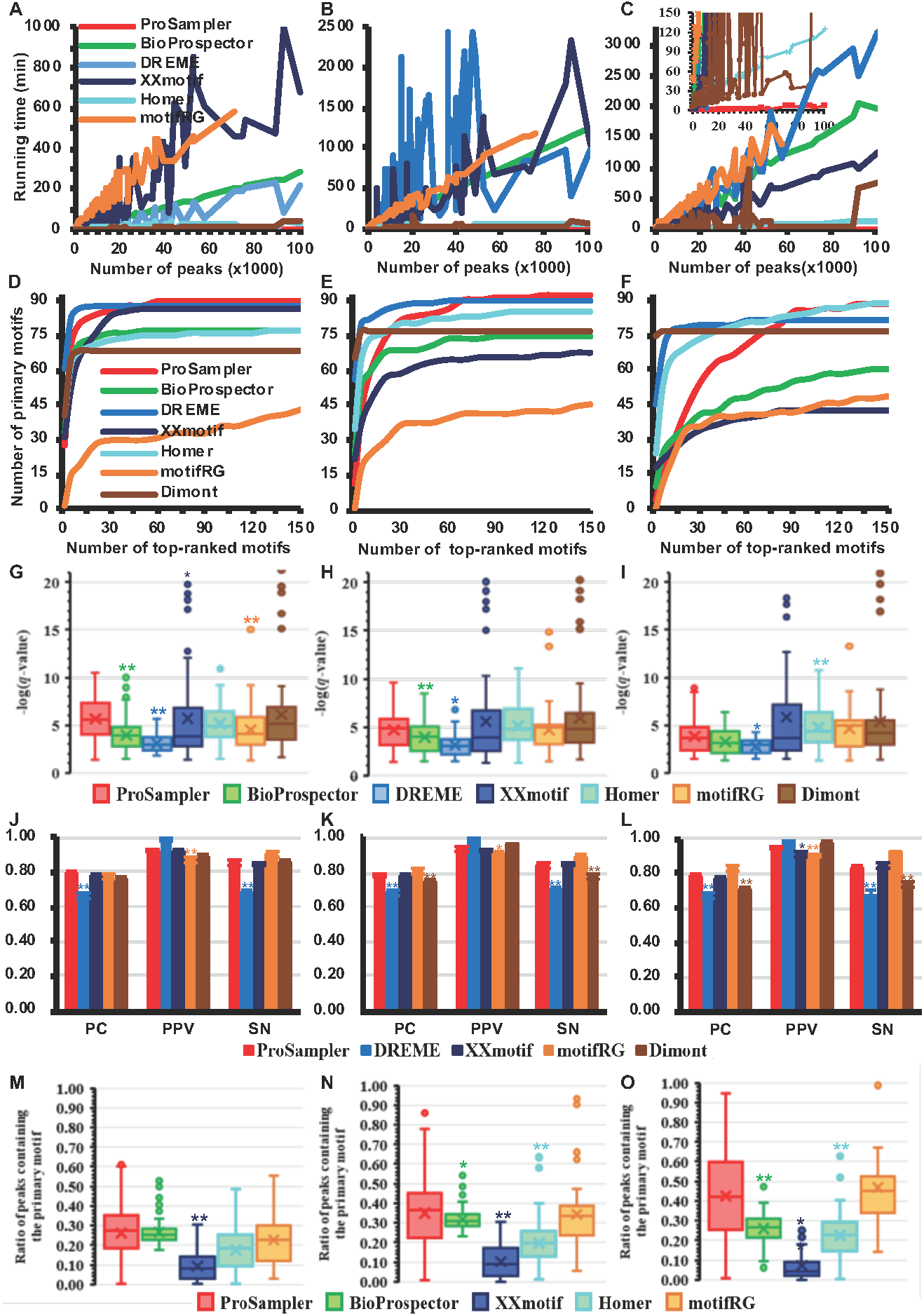
Comparison of the performance of the programs on the six synthetic datasets D_1_∼ D_6_ with various sizes. **A**. Running time of the six programs as a function of the number of sequences in the datasets. The inset is a blow-up view with a running time below 1,500 *s*. **B**. Number of recovered implanted motifs as a function of the number of top-ranked motifs predicted by the programs in the six datasets. **C**. Number of recovered implanted motifs as a function of the occurrence frequency of the implanted binding sites in the six datasets. **D.** Box-plot of the *q*-values of predicted motifs of the programs, matching the implanted motifs in the datasets by TOMTOM. * (for p<0.05) and (** for p<0.01) are Wilcoxon rank sum test significant levels between the result of the labeled program and that of ProSampler. **E**. Performance of the programs for predicting the lengths of implanted motifs in the datasets. * (for p<0.05) and (** for p<0.01) are t-test significant levels between the result of the labeled program and that of ProSampler. **F**. Average sensitivity of the programs for predicting the binding sites of implanted motifs as a function of the number of top-ranked motifs predicted by the programs in the six datasets. **G**. Average ROC curves of the programs for predicting the binding sites of implanted motifs in the six datasets. **H**. Average sensitivity of the programs for predicting the implanted binding sites as a function of their occurrence frequency in the six datasets. * (for p<0.05) and (** for p<0.01) are t-test significant levels between the result of the labeled program and that of ProSampler. The dotted line is the proportion of sequences found by ProSampler to contain the implanted binding sites.

We also compared ProSampler with three other programs DREME, motifRG and XXmotif for their ability to identify the length of implanted motifs in the synthetic datasets, using three metrics PC, PPV, and SN measuring the agreement between the lengths of implanted motifs and those of predicted matching motifs (see Online Methods for the definitions) in three perspectives. BioProspector and Homer were not included in this comparison as both are unable to automatically determine the lengths of motifs. As shown in Fig. 1E, ProSampler achieved the highest PC (0.94) and SN (0.96), both are significantly (p<0.05 or p<0.01, t-test) higher than those obtained by the other three programs except XXmotif for SN, for which the difference is not significant. ProSampler had comparable PPV (0.97) to the best PPV performer DREME (1.0), although ProSampler predicted the more implanted motifs. DREME had a perfect PPV of 1.0, as the most of implanted motifs had a length longer than 9 bp, while DREME was designed to identify motifs with a length ≤ 8 bp.

We have so far compared the accuracy of the tools at the motif level by comparing the returned motifs to the implanted ones. However, an ideal motif-finder should be able to identify all the binding sites of all motifs in a dataset, not just sub-motifs containing a subset of the binding sites of the motifs. To evaluate the programs in this regarding, we computed the sensitivity of a program for recovering the binding sites of the implanted motifs by its top-ranked motifs in the six datasets. DREME and Homer were not included in this evaluation as they only return the position weigh matrixes (PWMs) rather than the binding sites of predicted motifs. As shown in Fig.1F, ProSampler substantially outperformed the three other programs by identifying an average of 76.0% of the binding sites of the 60 implanted motifs by its 65 predicted motifs in the six datasets. Receiver operator characteristic (ROC) analyses indicate that ProSampler identified an average of 76.0% of the binding sites of the 60 implanted motifs in the six datasets with the lowest false positive rate of 0.005, substantially outperforming the three other programs (Fig.1G). We also evaluated the impacts of the occurrence frequency of implanted binding sites in a dataset on the ability of the programs to identify them. As shown in Fig. 1H, ProSampler found an average of 63.6%, 66.2%, 77.6%, 76.2%, 77.7%, 80.6%, 82.1%, 67.6%, 83.3% and 85.2% of the binding sites of motifs implanted with an occurrence frequency of 0.1, 0.2, 0.3, 0.4, 0.5, 0.6, 0.7, 0.8, 0.9 and 1.0 site/sequence, respectively, significantly (p<0.05 or p<0.01, t-test) outperforming the three other programs at all the occurrence frequencies except 0.3 and 0.6 site/sequence, at which Prosampler, Bioprospector and motifRG had similar performance (P>0.05). However, the Bioprospector (56) and motifRG (52) predicted fewer motifs than did ProSampler (60). As expected, the performance of ProSampler generally increased with the increase in the occurrence frequency of implanted motifs (Fig.1H). However, this was not the case for the other programs as their performance tended to decrease at higher frequencies (> 0.3 site/sequence) with large oscillations (Fig.1H). This is rather counter-intuitive, but the causes remain to be elucidated. Thus, ProSampler is more robust than the other programs to changes in the occurrence frequency of implanted motifs in the datasets.

In summary, ProSampler was orders of magnitude faster than the fastest existing tools on the synthetic datasets, and it substantially outperformed the existing tools in identifying the implanted motifs at both the motif and binding site levels in terms of sensitivity and specificity. ProSampler also had better than or comparable performance to the existing tools in determining the lengths of implanted motifs. Moreover, it was more robust than the existing tools to changes in the occurrence frequency of implanted binding sites in the synthetic datasets.

### Comparison of the programs for speed on real ChIP-seq datasets

We next compared ProSampler with six programs (BioProspector(Liu et al. 2001), DREME(Bailey 2011), XXmotif(Hartmann et al. 2013), Homer(S et al. 2010), motifRG(Yao et al. 2014) and Dimont(Grau et al. 2013)) on 105 real ChIP-seq datasets for 21 TFs, collected from embryonic stem (ES) cell-derived early human embryonic tissues using a micrococcal nuclease-based ChIP-seq (MNChIP-seq) technique(Tsankov et al. 2015). Each of these datasets contains 599∼ 100,778 binding peaks (Fig. S1) with an average length of 92∼ 1,151 bp (Fig. S2). To evaluate the effect of sequence lengths on the performance of the programs, we re-extracted binding peaks for each dataset with a length of 200, 500, and 1,000 bp centering on the summit of the originally called binding peaks (see Online Methods), forming three groups of 105 datasets: G_1_ (200 bp), G_2_ (500 bp) and G_3_ (1,000 bp). We let BioProspector output 150 motifs in each dataset, and the six other programs output all the motifs they found, but only considered up to the top 150 motifs in subsequent analyses. As shown in Figs. 2A ∼ 2C, ProSampler substantially outperformed in speed all the six other programs in the three groups of datasets (motifRG even crashed on some larger datasets in G_1_ ∼ G_3_, thus its running times on these datasets were not included). Specifically, ProSampler took only an average of 0.22, 0.54 and 1.17 min to finish on a dataset in the G_1_, G_2_ and G_3_ datasets, respectively, which on average was ∼ 85 fold faster than the runner-up Homer, ∼ 182 fold faster than XXmotif, and ∼ 264 fold faster than DREME. Notably, with the increase in the size of datasets, the running time of ProSampler increased largely linearly without much oscillation (Figs. 2A ∼ 2C), by contrast, those of the six other programs increased with large oscillations (Figs. 2A ∼ 2C) that were not seen on the synthetic datasets (Fig. 1A). These results suggest that the real datasets might be structurally more heterogeneous than the synthetic datasets, and the running times of the six other programs could be largely affected by the structures of the datasets, which had little effect on ProSampler’s running time. Therefore, ProSampler is not only orders of magnitude faster, but also more robust to the structures of datasets than the fastest existing tools.

**Fig. 2.**
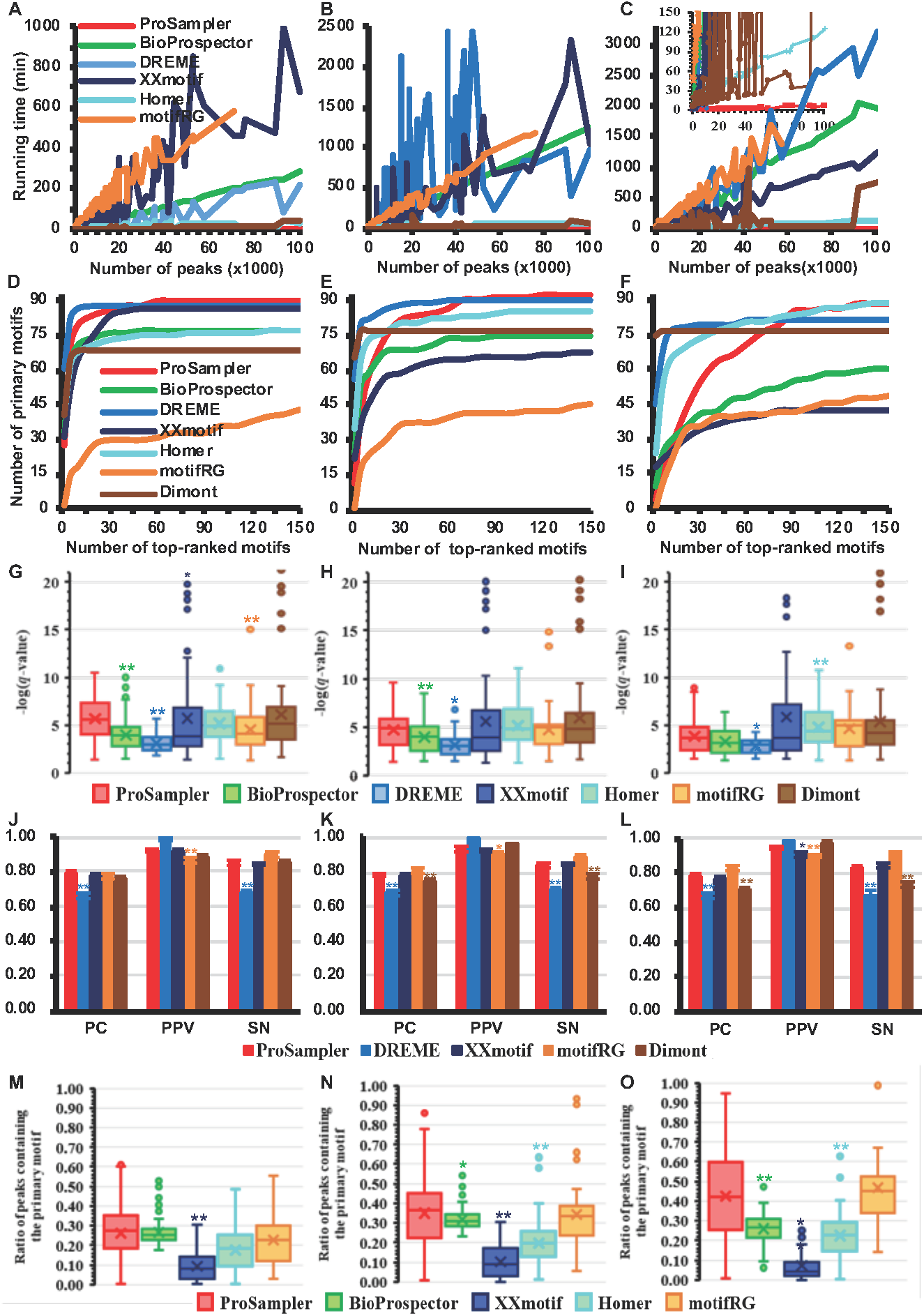
Performance comparison of the six programs for speed and identifying the primary motifs in real ChIP-seq datasets. **A, B** and **C**. Running time of the programs as a function of the size of the datasets in G_1_, G_2_ and G_3_, respectively. **D, E** and **F**. Cumulative number of primary motifs recovered by top-ranked motifs in the datasets in G_1_, G_2_ and G_3_, respectively. **G, H** and **I.** Box-plot of the *q*-values of predicted motifs of the programs, matching the primary motifs in the datasets in G_1_, G_2_ and G_3_, respectively, by TOMTOM. * (for p<0.05) and (** for p<0.01) are Wilcoxon rank sum test significant levels between the result of the labeled program and that of ProSampler. **J, K** and **L**. Performance of the programs for predicting the lengths of primary motifs in the datasets in G_1_, G_2_ and G_3_, respectively. * (for p<0.05) and (** for p<0.01) are t-test significant levels between the result of the labeled program and that of ProSampler. **M, N** and O. Proportion of sequences found by the programs to contain the binding sites of primary motifs in the datasets in G_1_, G_2_ and G_3_, respectively. *(for p<0.05) and (** for p<0.01) are t-test significant levels between the result of the labeled program and that of ProSampler.

### Comparison of the programs for identifying primary motifs in real ChIP-seq datasets

The major goal of a ChIP-seq experiment is to identify the primary motif of the ChIP-ed TF. To evaluate the programs for such capability, we counted the number of primary motifs recovered by each program in its top-ranked motifs in the 105 datasets in G_1_∼ G_3_. As shown in Fig. 2D and Table 2, ProSampler outperformed the six other programs in G_1_ by identifying 90 (85.7%) primary motifs, followed by DREME [88 (83.8%)], XXmotif [86 (81.9%)], BioProspector and Homer [77 (73.3%)], Dimont [68 (64.8%)] and motifRG [43 (41.0%)]. ProSampler also outperformed the six other programs in G_2_ by identifying 93 (88.6%) primary motifs in the 105 datasets, followed by DREME [90 (85.7%)], Homer [85 (81.0%)] and Dimont [77 (73.3%)], BioProspector [75 (71.4%)], XXmontif [68 (64.8%)] and motifRG [46 (43.8%)] (Fig. 2E and Table 2). Therefore, increase in the binding peak length from 200 bp (G_1_) to 500 bp (G_2_) increased the performance of ProSampler (90 vs 93), DREME (88 vs 90),Homer (77 vs 85)] and Dimont (68 vs 77) and motifRG (43 vs 46), presumably because the longer (500 bp) peaks include more binding sites of the target TFs than do the shorter (200 bp) peaks (see below), although most of the datasets have an average called binding peak length shorter than 500bp (Fig. S2). However, BioProspector (77 vs 75) and XXmontif (86 vs 68) had reduced performance on the 500 bp peaks. ProSampler was the runner-up in G_3_ by identifying 88 (83.8%) primary motifs in the 105 datasets, which is one fewer than the number that Homer [89 (84.8%)] found, followed by DREME [81(77.1%)], Dimont [77(73.3%)], BioProspector [60(57.1%)], motifRG [49(46.7%)] and XXmotif [42 (40.0%)] (Fig. 2F and Table 2). Therefore, further increase in the peak length from 500 bp (G_2_) to 1,000 bp (G_3_) reduced the performance of most programs including ProSampler (93 vs 88), DREME (90 vs 81), BioProspector (75 vs 60) and XXmotif (68 vs 42), but had no effect on Dimont (77 vs 77), and slightly increased the performance of Homer (85 vs 89) and motifRG (46 vs 49). Increase in the peak length to 1,500 bp reduced the performance of all the programs (data not shown), presumably because too long binding peaks might include more noise that interferes with motif finding.

**Table 2.** Comparison of performance of seven programs on all 105 MNChIP-seq datasets without running time limit.

| Programs | G1 (200 bp) |  |  |  | G2 (500 bp) |  |  |  | G3(1,000 bp) |  |  |  |
| --- | --- | --- | --- | --- | --- | --- | --- | --- | --- | --- | --- | --- |
|  | # of datasets with the PM found<sup>a</sup> | Ave. # of motifs found | Ave. # of known motifs<sup>b</sup> | Aver. # of cooperative motifs<sup>c</sup> | # of datasets with the PM found | Ave. # of motifs found | Ave. # of known motifs | Aver. # of cooperative motifs | # of datasets with the PM found | Ave. # of motifs found | Ave. # of known motifs | Aver. # of cooperative motifs |
| ProSampler | **90(85.7%)** | 49.6 | 12.9 | 2.7 | **93(88.6%)** | 93.8 | 17.9 | **3.4** | 88(83.8%) | 119.0 | **17.5** | **3.7** |
| BioProspector | 77(73.3%) | 150.0 | 8.7 | 0.9 | 75(71.4%) | 150.0 | 7.9 | 1.1 | 60(57.1%) | 150.0 | 5.9 | 0.8 |
| DREME | 88(83.8%) | 22.7 | 8.0 | 2.5 | 90(85.7%) | 42.1 | 9.0 | 2.4 | 81(77.1%) | 48.0 | 6.6 | 1.6 |
| XXmotif | 86(81.9%) | 49.4 | 12.4 | 2.7 | 68(64.8%) | 50.5 | 10.3 | 2.4 | 42(40.0%) | 54.3 | 6.4 | 1.5 |
| Homer | 77(73.3%) | 121.2 | **20.6** | **3.5** | 85(81.0%) | 136.9 | **18.9** | 3.2 | **89(84.8%)** | 147.1 | 17.0 | 2.8 |
| motifRG | 43(41.0%) | 126.8 | 9.4 | 1.8 | 46(43.8%) | 134.6 | 11.5 | 2.0 | 49(46.7%) | 132.5 | 13.1 | 2.2 |
| Dimont | 68(64.8%) | 4.3 | 1.6 | 0.6 | 77(73.3%) | 2.8 | 1.4 | 0.6 | 77(73.3%) | 2.0 | 1.3 | 0.5 |

<sup>a</sup> Number of datasets on which all the six programs successfully finished. <sup>b</sup> Numbers in bold type represent the best performance in the column. <sup>c</sup> Average number of predicted motifs matched to those in JASPAR in each dataset. <sup>d</sup> Average number of predicted motifs matched to those in TCoF-DB for the target TF in each dataset.

The motifs returned by ProSampler in the G_1_ datasets (Fig. 2G) also are significantly (p<0.05 or p<0.01) more similar to the known primary motifs than those found by BioProspector, DREME, XXmotif and mtifRG. Homer and Dimon have similar performance to ProSampler in this regard (p>0.05). In G_2_ ProSampler significantly (p<0.05 or p<0.01) outperformed BioProspector and DREME, but had similar performance to the remaining programs (p>0.05)(Fig. 2H). In G_3_ ProSampler significantly (p<0.05) outperformed DREME, but had similar performance to the remaining programs except Homer that outperformed ProSampler (p<0.01) (Fig. 2I). However, in all the three cases, ProSampler generally identify far more primary motifs (Fig. 2F and Table 2). The primary motifs identified by ProSampler in the 105 G_1_ datasets and their matched motifs in JASPAR are shown in Table 3, and those in 105 G_2_ and G_3_ datasets are shown in Tables S2 and S3, respectively.

**Table 3.**
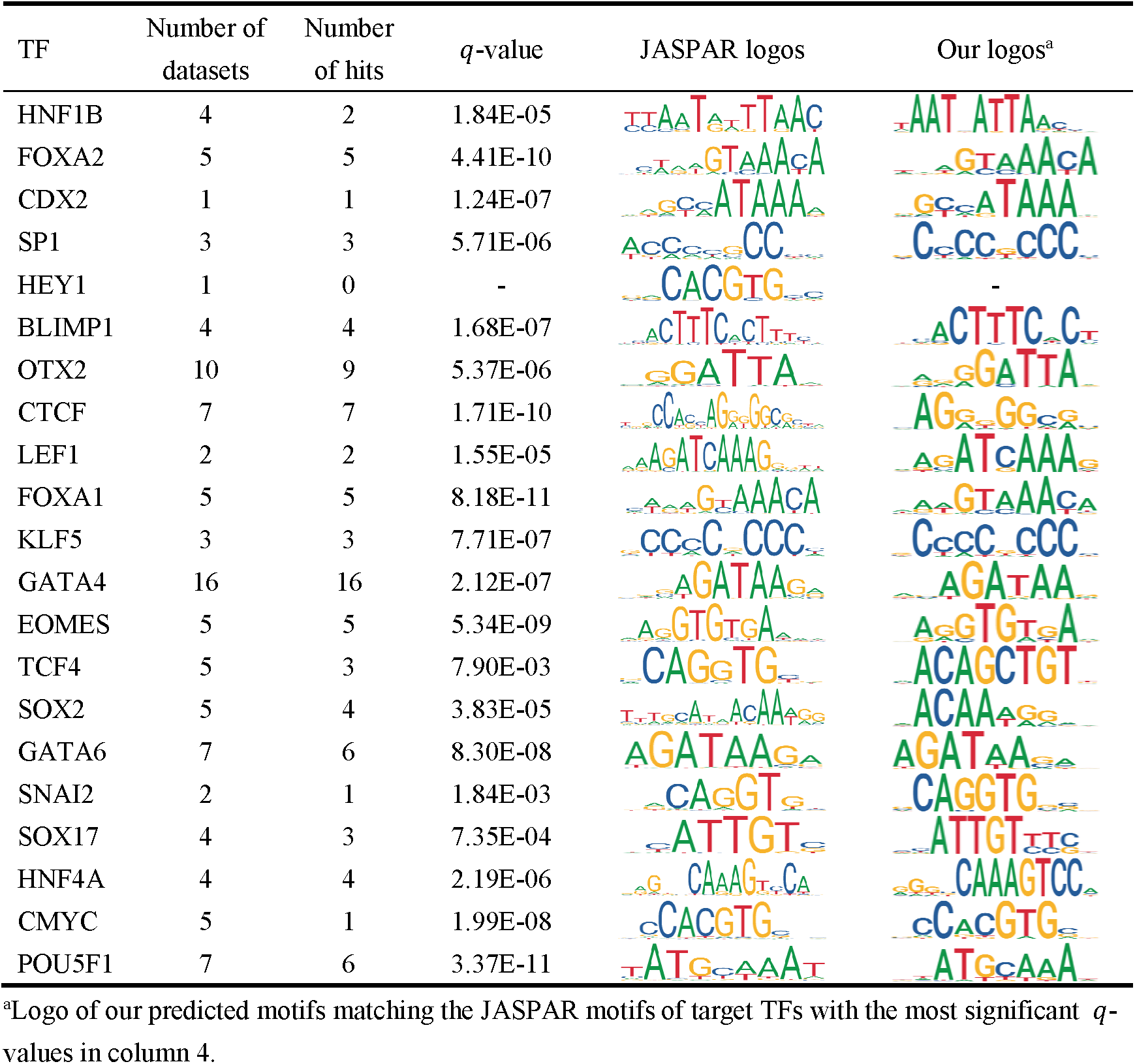
Motifs identified by ProSampler in the 105 ChIP-seq datasets of G1 (200 bp), matching JASPAR motifs of the target TFs.

We also compared ProSampler with XXmotif, DREME, motifRG and Dimont for identifying the lengths of primary motifs in the three groups of datasets. Since BioProspector and Homer cannot identify motif lengths by themselves, we did not include them in this evaluation. As shown in Fig.2J∼2L, ProSampler had significantly (p<0.05 or p<0.01) better performance than, or comparable performance to the other four programs for PC, PPV and SN.

Similar to the cases in the synthetic datasets, motifs predicted by different programs in the same dataset may match the target motifs significantly, but they may contain varying numbers of target binding sites located in varying numbers of peaks. A ChIP dataset is generally assumed to be enriched for binding peaks containing at least one binding site of the primary motif, thus a better motif-finder can find binding sites of the primary motif in more binding peaks than can a worse one. As shown in Figs. 2M∼2O, ProSampler generally identified significantly higher (p<0.05 or p<0.01, t-test) proportions of binding peaks in the datasets to contain putative binding sites of the primary motifs than the other programs (Figs. 2D∼2F). DREME, Homer and Dimont do not return binding site information, thus were not included in this analysis. However, the proportion of binding peaks found even by ProSampler to contain binding sites of primary motifs is surprisingly low with a median of 28%, 36% and 42% in G_1_, G_2_ and G_3_, respectively, and varies widely from as low as 0% to as high as 95% (Figs. 2M∼2O). In most datasets, this ratio was lower than the proportion of sequences found by ProSampler (48.3%) to contain the implanted binding sites in the synthetic datasets with a concentration of 0.6 site/sequence (the dotted line in Fig. 1H). These results suggest that on average more than 40% of “binding peaks” returned by a peak-calling algorithm might actually not contain the binding sites of the targeted TF, due probably to the low quality of the original data due to low specificity of the TF antibody used or other technical reasons. Alternatively, the target TF might bind these sequences indirectly through cooperative TFs, or the TF might bind other motifs in addition to the primary one.

### Comparison of the programs for identifying cooperative motifs in real ChIP-seq datasets

We designed the ProSampler algorithm to identify not only the primary motifs of the targeted TF, but also the motifs of cooperators in the binding peaks. Thus, we compared ProSampler with the six other programs for identifying cooperative motifs in the three groups of datasets. As shown in Figs. 3A∼3C and Table 2, in addition to the primary motifs, the programs returned a highly varying number of putative operative motifs in the three groups of 105 datasets. Specifically, Dimont found the smallest average number of 4.3, 2.8 and 2.0 putative motifs in G_1_, G_2_ and G_3_, respectively, followed by DREME (22.7, 42.1 and 48.0) and XXmotif (49.4, 50.5 and 54.3), while motifRG (126.8, 134.6 and 132.5),BioProspecter (150.0, 150.0 and 150.0) and Homer (121.2, 136.9 and 147.1) found the largest average number of putative motifs. Interestingly, ProSampler identified an intermediate average number of 49.6, 93.8 and 119.0 motifs in G_1_, G_2_ and G_3_, respectively, which are significantly (p<0.05 or p<0.01, Wilcoxon rank sum test) different from those obtained by the other programs except XXmotif in G_1_ (Figs. 3A∼3C). Since there is no golden standard benchmark to validate the prediction of cooperative motifs, we resorted to an alternative approach: we counted the number of known motifs as well as the number of known cooperative motifs of the primary motifs recovered by top-ranked motifs returned by each program. As shown in Fig. 3D and Table 2, in the G_1_ datasets, the cumulative number of known motifs recovered by the top-ranked motifs by both ProSampler and XXmotif increased rapidly and entered the saturation phase around the top 50 predicted motifs, close to the average number of predicted motifs in each dataset by both programs (Fig. 3A). ProSampler recovered a total of 1,357 known motifs in the G_1_ datasets, which is larger than those recovered by the other programs except Homer that recovered a total of 2,160 known motifs. In the G_2_ (Fig. 3E and Table 2) datasets, ProSampler (1,857) recovered approximately the same number of known motifs with that of Homer (1,968). Yet in the G_3_ datasets (Fig. 3F and Table 2), ProSampler outperformed all the six other algorithms by recovering the largest number of known motifs by most choices of N top-ranked motifs. Since the probability to find known motifs by chance is low, these matching motifs are likely to be true motifs, and may cooperate with the primary motifs in transcriptional regulation.

**Fig. 3.**
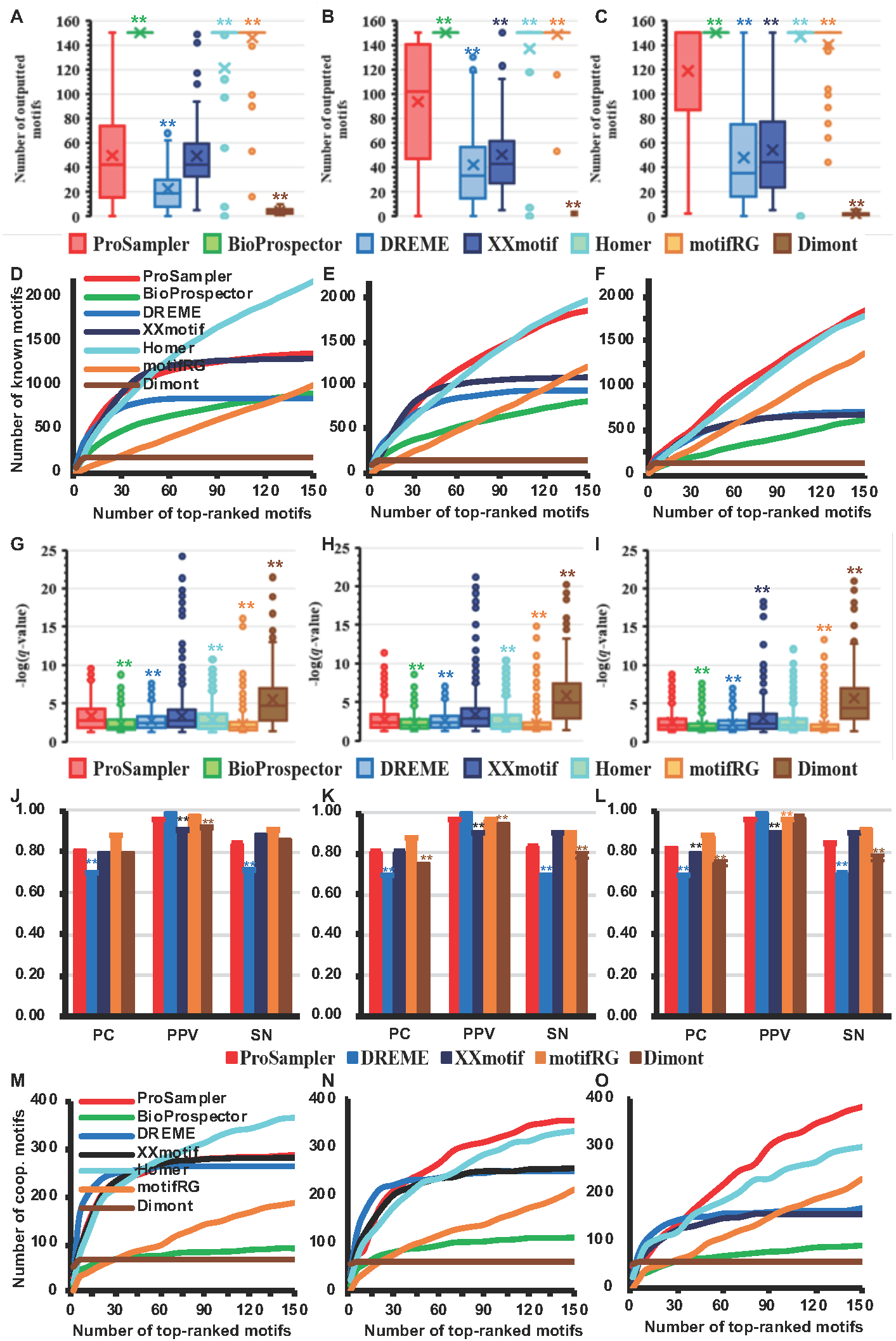
Performance comparison of the programs for identifying cooperative motifs in real ChIP-seq datasets. **A, B** and **C.** Number of motifs returned by the programs in the G_1_, G_2_ and G_3_ datasets, respectively. **D, E** and **F.** Cumulative number of known motifs matched by top ranked motifs of the programs in the G_1_, G_2_ and G_3_ datasets, respectively. **G, H, I**. Box-plot of the *q*-values of motifs predicted by the programs matching known motifs in the G_1_, G_2_ and G_3_ datasets, respectively, by TOMTOM. * (for p<0.05) and (** for p<0.01) are t-test significant levels between the result of the labeled program and that of ProSampler. **J, K** and *L*. Performance of the programs for predicting the lengths of known motifs in the G_1_, G_2_ and G_3_ datasets, respectively. * (for p<0.05) and (** for p<0.01) are t-test significant levels between the result of the labeled program and that of ProSampler. **M, N** and **O**. Cumulative number of the targeted TFs’ known cooperative motifs matched by top ranked motifs of the programs in the G_1_, G_2_ and G_3_ datasets, respectively.

Moreover, as shown in Figs. 3G∼3I and Table 2, the motifs returned by ProSampler in all the three groups of datasets are significantly (p<0.05 or p<0.01) more similar to the known motifs than those predicted by all the other programs except that XXmotif performed significantly (p<0.01) better in G_3_ and Dimont in G_1_∼ G_3_ (p<0.01). However, as a tradeoff, both XXmotif and Dimont predicted far fewer known motifs (Figs. 3D∼3F). For identifying the lengths of the matched known motifs in the three groups of datasets, ProSampler had significantly (p<0.01) better performance than, or comparable performance to the other four programs for PC, PPV and SN (Figs. 3J∼3L). The results are largely similar to those obtained for the primary motifs (Figs. 2J∼2L).

ProSampler also identified the largest average number (3.4 and 3.7) of motifs in G_2_ and G_3_ matching those of known cooperative TFs of the target TFs documented in the TcoF-DB database (Fig. 3M∼3O and Table 2)(Schaefer et al. 2011; Schmeier et al. 2017) (see Table S4 for all known cooperative factors of the 21 target TFs), although Homer (3.5) outperformed ProSampler (2.7) in G_1_. These matching motifs are likely to be true cooperative motifs of the primary ones. As expected, with the increase in the binding peak length of the datasets, ProSampler identified an increasing average number of known cooperative motifs. However, the reverse, an unexpected resul, was true for Homer for unknown reasons.

### Patterns of motif occurrence in ChIP-seq datasets

After demonstrated the superiority of ProSampler for identifying both primary as well as cooperative motifs in even very large ChIP-seq datasets, we asked what the patterns are of the occurrence of predicted binding sites in the binding peaks, and why an average of more than half of the binding peaks lack the predicted primary motifs (Figs. 2M∼2O), and why the primary motifs could not be found in some datasets. To this end, we split each dataset in two subsets S_P_ and S_N_, where S_P_ consists of the binding peaks in which at least one binding site of the primary motif was predicted, and S_N_ consists of those in which no binding site of the primary motif was found. Clearly, in the datasets where primary motifs were not found, S_P_ is empty. We aligned all the sequences in each subset with the summits of binding peaks being the center (with coordinator 0), and for each nucleotide position along the alignment, we counted the number of predicted binding sites overlapping the position over all the binding peaks in the subset. The results from the G_2_ datasets are shown in the Supplementary Data File 1, and will be used in the following analyses. Similar results were found in the G_1_ and G_3_ datasets (data not shown).

We note that the distribution of putative binding sites along the binding peaks in the two subsets can be classified in four patterns (Supplementary Data File 1): 1) in the “1 0” pattern, the predicted binding sites have a bell-shape distribution around the summits of the binding peaks in S_P_, but a largely uniform distribution in S_N_, indicating enrichment of predicted binding sites around the summits of the binding peaks in S_P_ but not in S_N_. There are 59 (56.2%) of the 105 datasets in G_2_ with this pattern of distribution (Supplementary Data File 1). Figs. 4A and 4B show the cases for the datasets GSM1505766 of SOX2 and GMS1505769 for SP1, respectively. 2) In the “1 1” pattern, the predicted binding sites have a bell-shaped distribution around the summits of the binding peaks in both S_P_ and S_N_, indicating enrichment of predicted binding sites around the summits of the binding peaks in both S_P_ and in S_N_. There are 22 (21.0%) datasets in G_2_ having this pattern of distribution (Supplementary Data File 1). Figs. 4C and 4D shows the cases for the datasets GSM1505618 of BLIMP1/PRDM1 and GSM1505636 of FOXA1, respectively. 3) In the “0 1” pattern, the predicted binding sites have a largely uniform distribution in S_P_, or S_P_ is empty, but a bell-shaped distribution in S_N_, indicating enrichment of predicted binding sites around the summits of the binding peaks in S_N_, but not in S_P_. There are 10 (9.5%) datasets in G_2_ having this pattern of distribution (Supplementary Data File 1).Figs. 5A and 5B shows the cases for the datasets GSM1505789 for TCF4 and GSM1505760 of SOX17, respectively. And 4) in the “0 0” pattern, the predicted binding sites have a largely uniform distribution in both S_P_ and S_N,_ or S_P_ is empty, indicating no enrichment of predicted binding sites around the summits of the binding peaks in both S_P_ and S_N_. There are 14 (13.3%) datasets in G_2_ with this pattern of distribution (Supplementary Data File 1), Figs. 5C and 5D shows the cases for the datasets GSM1505759 of SNAI2 and GSM1505682 of HNF1B, respectively.

**Fig. 4.**
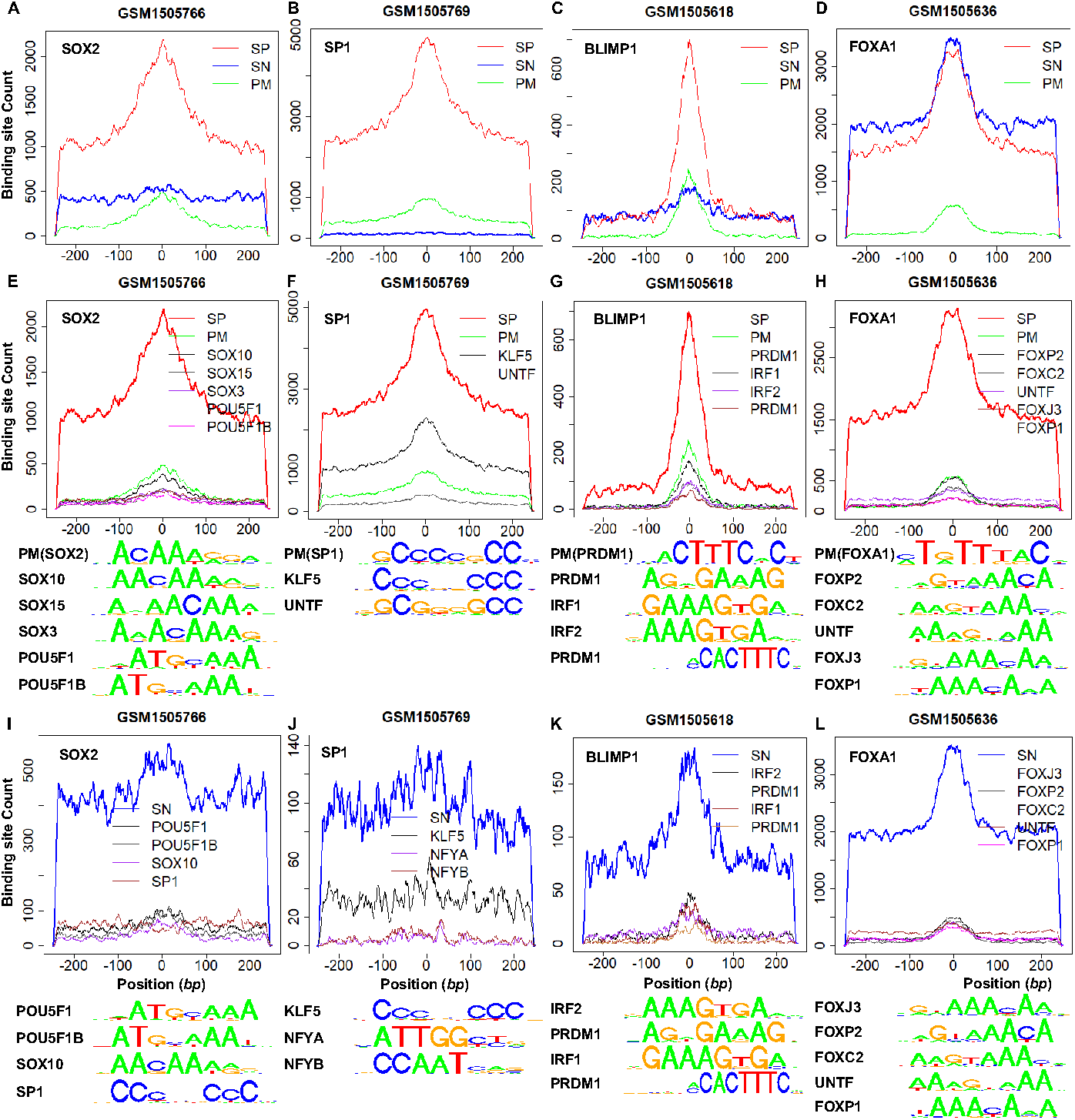
Examples of the “1 0” and “1 1” distribution patterns of predicted binding sites in the S_P_ and S_N_ subsets. **A, B, C and D**. Distributions of the predicted binding sites in the S_P_ and S_N_ subsets of datasets GSM1505766 of SOX2, GSM1505769 of SP1, GSM1505618 of BLIMP1 and GSM1505636 of FOXA1, respectively. The distributions of the binding sites of the primary motifs (PM) are also shown. **E, F, G and H**. Distributions of the most dominant motifs and their logos in the S_P_ subsets of datasets GSM1505766, GSM1505769, GSM1505766 and GSM1505769, respectively. **I, J, K and L**.Distributions of the most dominant motifs and their logos in the S_N_ subsets of datasets GSM1505766, GSM1505769, GSM1505766 and GSM1505769, respectively.

**Fig. 5.**
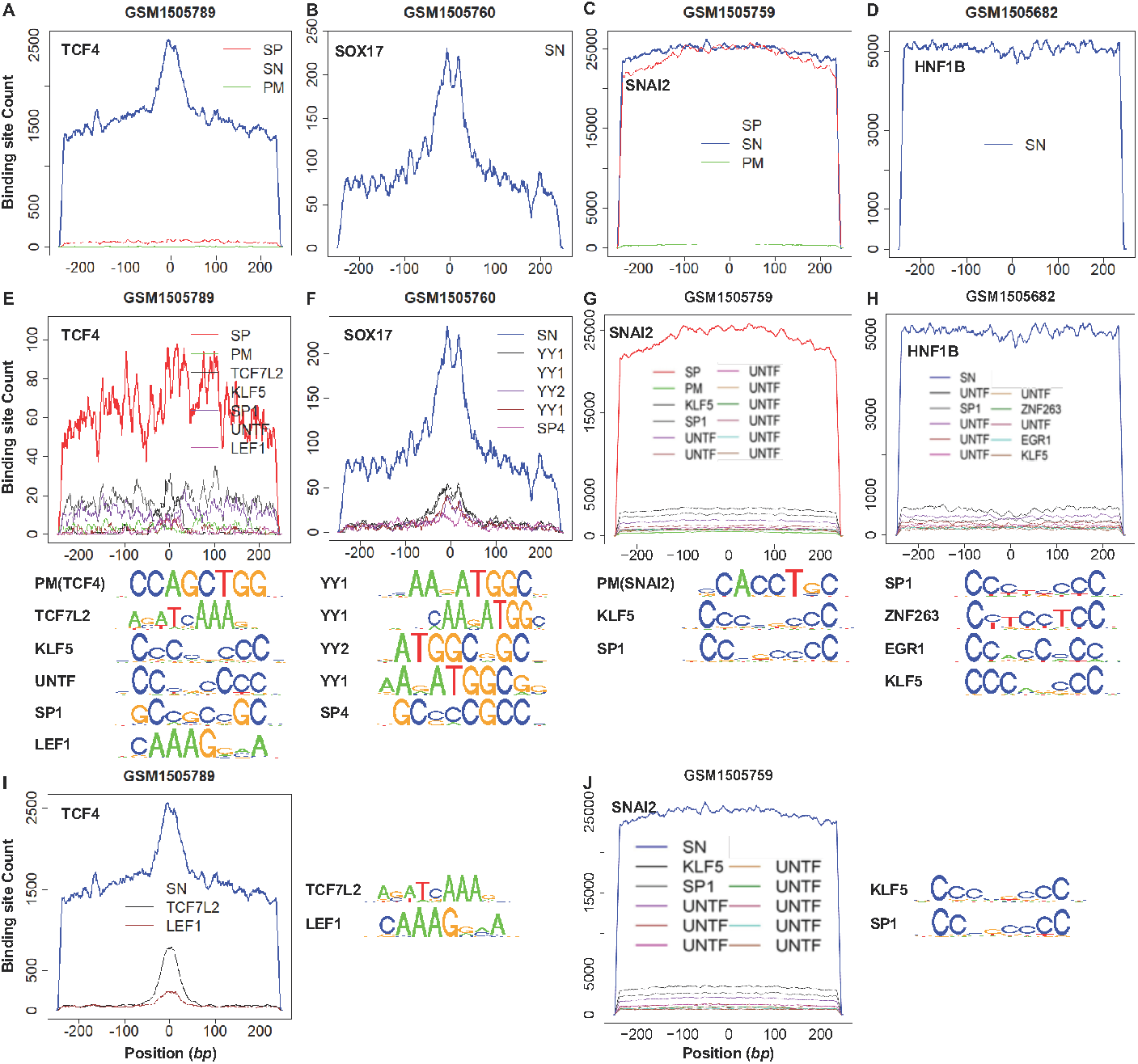
Examples of the “0 1” “and”0 0“ distribution patterns of predicted binding sites in the S_P_ and S_N_ subsets. **A, B, C and D**. Distributions of the predicted binding sites in the S_P_ and S_N_ subsets of datasets GSM1505789 of TCF4, GSM1505760 of SOX17, GSM1505759 of SNAI2 and GSM1505682 of HNF1B, respectively. The distribution of binding sites of the primary motifs (PM) in GSM1505789 and GSM1505759 are also shown. Primary motifs were not found in GSM1505760 and GSM1505682, so their S_P_s are empty. **E**, **and G**. Distributions of the most dominant motifs and their logos in the S_P_ subsets of datasets GSM1505789 and GSM1505682, respectively. **F, H, I and J**. Distributions of the most dominant motifs and their logos in the S_N_ subsets of datasets GSM1505760, GSM1505682, GSM1505789 and GSM1505759, respectively.

As expected, in most (81 or 87.1%) of the 93 datasets in which primary motifs were found, the predicted binding sites of the primary motif in the S_p_ subsets also had a bell-shaped distribution around the summits of binding peaks (Figs. 4A and 4B as examples for the “1 0” pattern, and 4C and 4D as examples for the “1 1” pattern, see Supplementary Data File 1 for other datasets), these putative binding sites of the primary motifs might lead to the pulldown of these sequences. Interestingly, the bell-shaped distributions of the predicted binding sites of the primary motifs in these 81 datasets are much lower than those of all the predicted binding sites in the S_P_ sub-datasets (Figs. 4A∼4D, see Supplementary Data File 1 for other datasets), indicating that some non-primary motifs are also enriched around the summits of binding peaks in the S_p_ subsets, and they are likely to be cooperative motifs of the primary motifs. Indeed, some of these enriched motifs match known cooperative motifs of the primary motifs, while the remaining ones might be novel cooperative motifs of known and unknown cooperators. For instance, in the dataset GSM1505766 of SOX2, in addition to the primary motif of SOX2, known motifs of SOX3, SOX10, SOX15, POU5F1/OCT-4 and POU5F1B have a bell-shaped distribution, largely accounting for the distribution of binding sites in the S_P_ subset (Fig. 4E). It has been shown that SOX2 and SOX3 cooperatively regulate otic/epibranchial placode induction in zebrafish(Gou et al. 2018), and SOX2 and SOX10 can physically interact with positive transcription elongation factor b in Schwann cells(Arter and Wegner 2015). SOX2 and POU5F1/OCT4 work as transcriptional activators in reprogramming human fibroblasts(Narayan et al. 2017), and SOX2 is transcriptionally regulated by an enhancer containing a composite SOX-OCT element that POU5F1 and SOX2 bind in a combinatorial interaction(Chew et al. 2005). In the case of the dataset GSM1505769 of SP1, in addition to the primary motif, motifs of KLF5 and an unknown TF (UNTF) with a bell-shaped distribution are largely responsible for the bell-shaped distribution of putative binding sites in S_P_ (Fig. 4F). It is known that SP1 and KLF5 cooperatively regulate Am80-induced apelin expression through their direct binding to the apelin promoter(Lv et al. 2013). As the motif of the unknown TF is quite different from that of SP1 (Fig. 4F), this unknown TF might physically interact with SP1, leading to the pulldown of the relevant sequences. In the dataset GSM1505618 of BLIMP1/PRDM1, in addition to the primary motif (MA0508.1), two variants of the PRDM1 motif, and motifs of IRF1 and IRF2 were enriched around the summits of binding peaks, accounting for the bell-shaped distribution of the putative binding sites in the S_P_ subset (Fig. 4G). It has been reported that the IRF-element (IRF-E) bound by IRF1 and IRF2, can also be bound by BLIMP1/PRDM1 *in vitro*(Tooze et al. 2006), probably due to their high similarity (Fig. 4G). In the dataset GSM1505636 of FOXA1, in addition to the primary motif, similar motifs of four FOX TFs (FOXP1, FOXP2, FOXC2, and FOXJ3) as well as a very different motif of an unknown TF were enriched around the summit of binding peaks, largely accounting for the bell-shaped distribution of the predicted binding sites in the S_P_ subset (Fig. 4H). It has been shown that FOXA1 co-expresses with FOXP1 and FOXC2 during adipogenesis(Gerin et al. 2009) and may have physical interactions with them(Hu et al. 2015).

However, in the remaining 12 (mainly for CTCF, LEF1, SNAI2, and TCF4, Supplementary Data File 1) of the 93 datasets where primary motifs were found, the predicted binding sites of the primary motifs in the S_p_ subsets are largely uniformly distributed along the binding peaks (Figs. 5A as an example for the “0 1” pattern and 5C as an example for the “0 0” pattern), indicating either low quality of the datasets or a broad distribution of the binding sites of primary motifs along the binding peaks. In these 12 datasets, the count of all the predicted binding sites in the S_P_ sub-datasets also has a largely uniform distribution along the binding peaks (Supplementary Data File 1). The count could be slightly higher than that of the primary motif as in the case of GSM1505789 of TCF4 (Fig. 5A), indicating few binding sites of non-primary motifs found in the S_P_ subset. The count could also be much higher than that of the primary motif as in the case of GSM1505759 of SNAI2 (Fig. 5C), indicating a considerable number of binding sites of non-primary motifs found in the S_P_ subset. The most dominant non-primary motifs in the S_P_ subsets all show a largely uniform distribution alone the binding peaks (Figs. 5E and 5G). In the case of GSM1505789 of TCF4, a few most dominant non-primary motifs largely account for the distribution of all the predicted binding sites in the S_P_ dataset (Fig. 5E), while this is not true in the case of GSM1505759 of SNAI2 (Fig. 5G). Only few of these motifs match those of known cooperators of the target motifs (such as LEF1 for TCF4 in GSM1505789), therefore, although they are still likely to be cooperative motifs of the primary motifs, more caution should be taken for this conclusion.

As we indicate earlier, the S_N_ subsets make up an average of 65% of the datasets (Fig. 2N).Nonetheless, the failure to find primary motifs in the S_N_ sub-datasets cannot be fully explained by the possible inability of ProSampler to identify the binding sites, because ProSampler can only misses ∼ 15% binding sites in datasets with a motif occurrence frequency of 1 site/sequence according to the results from the synthetic datasets (Fig. 1H). To reveal the possible underlying causes, we noted that, in most (73 or 69.5%) of the 105 datasets, the predicted binding sites in the S_N_ sub-datasets have a largely uniform distribution (Figs. 4A and 4B as examples for the “1 0” pattern, and 5C and 5D as examples for the “0 0” pattern, see Supplementary Data File 1 for other datasets), and the same is largely true for the most dominant motifs found in these S_N_ subsets (Figs. 4I, 4J, 5J and 5H), indicating little or no enrichment of predicted binding sites around the summit of the binding peaks in these S_N_ datasets as expected. In the S_N_ subsets of GSM1505766 of SOX2 and GSM1505769 of SP1 with the “1 0” pattern, motifs of known cooperators of the target TFs were found, such as POU5F1, POU5F1B and SOX10 for SOX2 (Fig. 4I), and KLF4, NFYA and NFYB for SP1(Figs 4J). Thus, the sequences in these S_N_ subsets might be pulldown indirectly by the physical interaction of the target TFs with their cooperators. In the S_N_ subsets of GSM1505759 of SNAI2 (Fig. 5J) and GSM1505682 of HNF1B (Fig. 5H) with the “0 0” pattern, although some of the dominant motif match known motifs, none of them match those of the known cooperators, or resembles the primary motifs. Therefore, we argue that at least most of the sequences in these S_N_ subsets might not contain binding sites of primary motifs, and they might be mistakenly called as “binding peaks”, probably due to low quality of datasets for various technical reasons.

On the other hand, in 32 (37.1%) of the 105 datasets, the predicted binding sites in the S_N_ subsets have a bell-shaped distribution around the summits of the binding peaks (Fig. 4C and 4D as examples for the “1 1” pattern, and Fig. 5A and 5B as examples for the “0 1” pattern, see Supplementary Data File 1 for other datasets), indicating that putative binding sites of non-primary motifs were somehow enriched around the summits of the binding peaks. In the datasets GSM1505618 of BLIMP1/ PRDM1 and GSM1505636 of FOXA1 with the “1 1” pattern, except for the primary motif, the motifs of the same TFs enriched in the S_P_ subsets also were enriched around the summit of binding peaks in the S_N_ subset, such as PRDM1, IRF1 and IRF2 in the former case (Fig 4K), and FOXC2, FOXJ3, FOXP1, POXP2 and an UNTF in the latter case (Fig. 4L). The distributions of these motifs largely account for the bell-shaped distribution of the putative binding sites in the S_N_ subsets (Figs. 4K and 4L). Therefore, possible crossing binding of the target TFs to these highly similar binding motifs(Tooze et al. 2006) (Figs. 4K and 4L), and/or physical interactions between the target TFs and the cooperators(Hu et al.2015) might lead to the pulldown of the sequences in the S_N_ subsets. In the dataset GSM1505789 of TCF4 with the “0 1” pattern, where the primary motif was found to have a largely uniform distribution (Figs. 5A), motifs of TCF7L2 and LEF1 were enriched around the summit of the binding peaks in the S_N_ subset, accounting for the bell-shaped distribution of the putative binding sites (Fig. 5I). Although TCF4, TCF7L4 and LEF1 belong to the same TF family (Hrckulak et al. 2016), the motif of the target TF TCF4 is quite different from those of TCF7L4 and LEF1 (Figs. 5E and 5I), thus TFC4 is less likely to bind the motifs of the latter two TFs. Nonetheless, LEF1 is a cooperator of TCF4(Schaefer et al.2011; Schmeier et al. 2017), therefore, it is likely that physical interaction between TCF4 and LEF1 might result in the pulldown of the sequences in the S_N_ subset. In the dataset GSM1505760 of SOX17 with the “0 1” pattern, where the primary motif of the target TF was not found (Fig. 5B), three variants of the YY1 motif as well as motifs of YY2 and SP4 were enriched around the summit of the binding peaks in the S_N_ subset (Fig. 5F). Since the motif of the target TF SOX17 is very different from those of YY1, YY2 and SP4 (Fig. 4F and Table S2), it is unlikely that TFC4 might bind them. Furthermore, SOX17, SP4 and the YY proteins belong to different protein families, excluding the possible crossing binding of the antibody against SOX17 used in the experiment. Therefore, these sequences might be pulled down indirectly through physical interactions between SOX17 and these proteins.

### Predicting known and novel cooperative motifs

An average of 20.1 known cooperative TFs have been documented in TcoF-DB(Schaefer et al. 2011; Schmeier et al. 2017) for the 21 target TFs (Table S4), ranging from 0 (for EOMES) to 103 (for CMYC). For the 20 target TFs with at least one known cooperative TF, ProSampler identified motifs of an average of 5.7 (27.0%) of the known cooperative TFs (Table 4). As expected, motifs of some targeted TFs were identified as one of cooperative motifs in the datasets for their known cooperative TFs, such as SOX2 and POU5F1, TCF4 and LEF1, and OTX2 and FOXA2, to name a few (for more see Table 4). Moreover, we predicted an average of 14.5 motifs/dataset in the 105 G_2_ datasets (Table 2), which match known motifs in JASPAR but not those of TFs in TcoF-DB (Table S4). Pooling all these matching motifs from all the datasets for each target TF, we identified from 36 (for HEY1) to 387 (for GATA4) (an average of 218.0) known motifs in the datasets for each target TF (Fig. 6A). Due to their high similarity to known motifs and enrichment around the summit of the binding peaks, their cognate TFs are likely to be novel cooperators of the target TFs in transcriptional regulation (Table S5). Additionally, there are an average of 75.9 predicted motifs that do not match any known motifs in JASPAR. Many of them are highly similar to one another and can be clustered into 640 clusters based on their similarity(Zhang et al. 2013) using the CLIMP program(Zhang and Chen 2016), with vast majority (71.1%) clusters containing more than two motifs (Fig. 6B). We consider each of these 640 motif clusters a unique motif whose distribution for the 21 target TFs are shown in Fig. 6C and logos are listed in the Supplementary Data File 2. As highly similar spurious motifs are unlikely to be found by chance in multiple datasets, at least the majority of these unique motifs are likely to be novel cooperative motifs whose cognate TFs are yet to be identified.

**Table 4.** Predicted motifs in the 105 MNChIP-seq datasets of G_2_ (500 bp) matching known cooperative motifs of the ChIP-ed TFs.

| TF | Known co-factors | Discovered cooperative motifs | Cooperators |
| --- | --- | --- | --- |
| CTCF | 10 | 2 | TFAP4, PKNOX1 |
| LEF1 | 13 | 5 | MITF, EGR1, MYB, TCF4, TCF7L2 |
| FOXA2 | 13 | 3 | GSC, OTX2, EN2 |
| SOX2 | 86 | 15 | TWIST1, SOX5, RFX1, SOX6, SP2, KLF4, ARID3B, TCF3, ZEB1, ALX4, TCF12, ARID3A, JUND, RFX3, POU5F1 |
| CMYC | 103 | 20 | ELF3, ZBTB18, JUN, ETV3, MAX, YY1, HIF1A, FOSL1, TCF12, NFYB, NMYC, FOSL2, ETV6, NFE2L2, ZNF740, MNT, NFIL3, SP1, ESR1, FOXP1 |
| KLF5 | 16 | 2 | SREBF1, JUN |
| GATA4 | 18 | 5 | ALX4, SP1, ZBTB3, ID3, ID2 |
| BLIMP1 | 1 | 0 | - |
| CDX2 | 3 | 0 | - |
| GATA6 | 5 | 1 | SP1 |
| HNF1B | 3 | 3 | ATF1, HNF1A, CREB1 |
| OTX2 | 1 | 1 | FOXA2 |
| SOX17 | 2 | 1 | TCF7L2 |
| FOXA1 | 14 | 5 | TFAP4, GATA3, NFIC, CREB1, AR |
| SNAI2 | 1 | 0 | - |
| SP1 | 66 | 29 | TFAP4, ARNT, SREBF2, JUN, YY1, ETS1, E2F1, E2F3, EGR1, NFYB, NMYC, SP3, RXRA, NFYA, NR2F1, ELF1, KLF4, ZBTB7A, POU2F2, AR, GABPA, SREBF1, POU2F1, E2F2, MYOD1, ESR1, STAT3, MYOG, CMYC |
| HEY1 | 7 | 0 | - |
| POU5F1 | 21 | 4 | FOXD3, SOX2, KLF4, PBX1 |
| TCF4 | 19 | 14 | LEF1, TAL1, TWIST2, PPARG, ETS1, TCF12, BHLHA15, EGR1, MYOD1, ID2, MYOG, TCF21, ID4, MSC |
| HNF4A | 21 | 4 | SREBF1, SP1, SREBF2, NR2C2 |

**Fig. 6.**
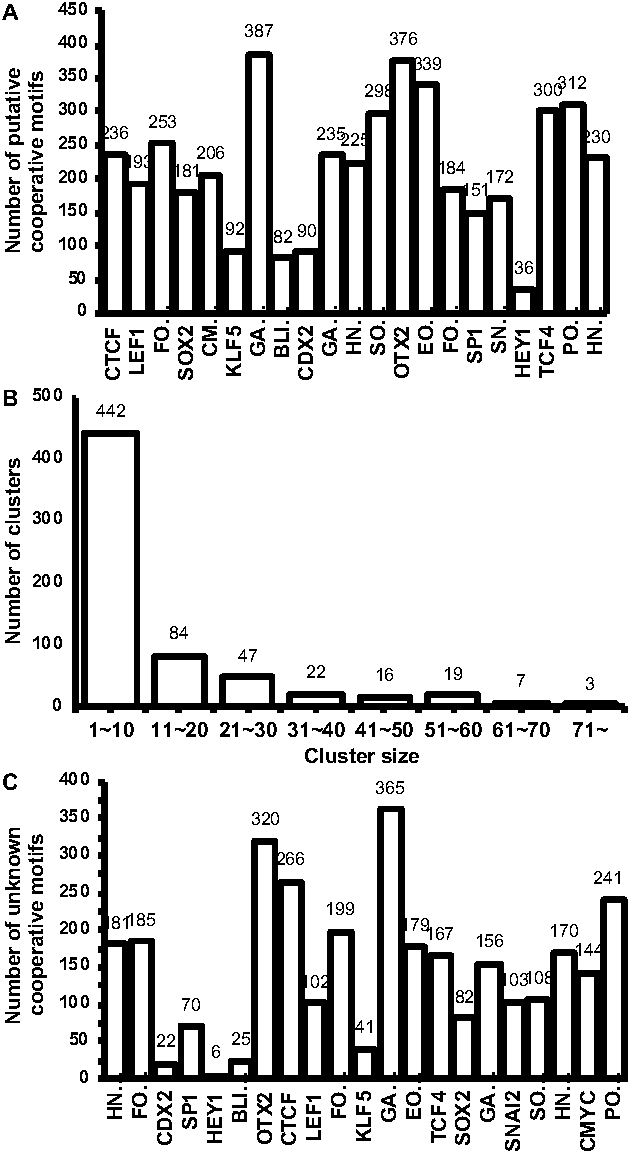
Prediction of cooperative motifs for the 21 target TFs based on the 105 ChIP-seq datasets (G_2_).**A**. Number of predicted cooperative motifs for the target TFs, which match known motifs in JASPAR but those of TFs in TcoF-DB. **B**. Distribution of the size of the motif clusters composed of motifs that do not match known motifs in JASPAR. **C**. Number of predicted cooperative motifs for the target TFs, which do not match known motifs in JASPAR.

## DISCUSSION

We designed ProSampler aiming at finding not only the primary motifs of the targeted TFs, but also motifs of cooperators in the binding peaks with a length of typical CRMs (500∼ 1,000 bp) in very large ChIP-seq dataset. To this end, we took the following tactics: 1) Instead of performing Gibbs sampling on original sequences directly(Liu et al. 2001; Frith et al. 2008), we sample on preliminary motifs formed by combining highly similar significant *k*-mers, with the aid of the motif similarity graph. As the number of possible *k*-mers is fixed in any size of data, this step runs in an almost constant time. 2) Unlike most of the existing algorithms that identify the motif length by exhaustively evaluating each length within a specified interval (Bailey and Elkan 1994; Bailey et al. 2010; Bailey 2011), we determine the motif length by extending the *k*-mer core motif using a two-proportion *z*-test, saving a few fold of CPU time. 3) By storing the flanking *l*-mers in memory, we avoid extensive I/O. 4) We combine the strength of *k*-mer numeration and Gibbs sampling approaches to identify subtle weak motifs. Indeed, as we demonstrated in this work, these strategies render ProSampler to outperform the six state-of-the-art tools in speed and accuracy in identifying the primary motifs as well as cooperative ones in very big ChIP-seq datasets with a length of typical CRMs. Thus, ProSampler allows researchers to fully exploit valuable ChIP-seq datasets and identify all possible TFBSs enriched in them. The results can provide new insights into the cooperative regulation of gene transcription by multiple TFs and possible technical issues in generating the datasets.

Indeed, using ProSampler, we found four patterns of binding sites occurrence along the binding peaks in the 105 MNChIP-seq datasets evaluated. These patterns can be well explained by the identified motifs in the datasets, and thus the results can throw light on the mechanisms of cooperative TF binding in CRMs, or pinpoint possible technical problems in data generation. For instance, these patterns can explain why a considerable proportion of binding peaks in a dataset do not harbor a binding site of the target TF: in a dataset with the “1 1” or “0 1” pattern, the sequences in its S_N_ subset with no binding site of the primary motif found contains binding sites of the cooperators of the target TF in the summit of the binding peak (Figs. 4K, 4L, 5F and 5I), thus might be pull down indirectly through physical interaction between the target TF and the cooperators. In a dataset with a “1, 0” pattern, the sequences in its S_N_ subset contain binding sites of the cooperators of the target TF anywhere along the binding peak (Figs. 4I and 4J), thus might be pull down indirectly through more complex physical interactions between the target TF and the cooperators. In a dataset with a “0 0” pattern, motifs found in its S_N_ subset usually do not match those of the known cooperators or resemble the primary motifs (Figs. 5H and 5J), thus they might be mistakenly called as “binding peaks”, probably due to low quality of datasets for various technical reasons. We also predicted an average of 218.0 known (Fig. 6A) and 75.9 unknown (Fig. 6B) motifs as novel cooperative motifs for each of the 21 target TFs using the 105 datasets and identified a total of 640 putative novel motifs of unknown TFs. In this regarding, ProSampler can be used to predict more cooperative motifs in an even larger number of ChIP-seq datasets, and the results can be integrated to predict all CRMs in genomes as recently demonstrated(Niu et al. 2018).

## METHOD

### Synthetic datasets

We downloaded the known vertebrate TF binding motifs (pfmVertebrates.txt) and the background sequences (upstream1000.fa) from the JASPAR database(Portalescasamar et al. 2010). The pfmVertebrates.txt file contains 519 motifs with a length ranging from 4 to 21 bp. The upsteam1000.fa file contains 43,632 upstream regions of genes with a length of 1,000 bp. To generate six synthetic datasets D_1_∼ D_6_, we first randomly selected 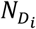 (500, 1,000, 2,000, 5,000, 10,000 and 20,000) sequences from upstream1000.fa and shuffled the sequences in each dataset. Then for each dataset *D*_*i*_ containing 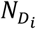 sequences, we randomly chose ten motifs *M*_1_, *M*_2_, …, *M*_10_ from pfmVertebrates.txt for distinct TF families, with a length *l* ranging from 6 to 15 bp, and implanted them at a frequency *α*= 0.1, 0.2, …, 1.0 site/sequence, respectively, in the 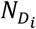 sequences in the dataset. Specifically, for each selected motif, we randomly chose 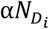 sequences with replacement from the dataset, randomly selected a position in a sequence chosen, and substituted the subsequence starting at this position with a DNA sequence generated according to the motif’s position frequency matrix (PFM) within a Hamming Distance cutoff to the consensus string of this PFM. We recorded the substituted positions and avoided overlapping implanting. Note that with replacement sampling, we allow the ZOOPS (zero-or-one occurrence per sequence) motif distribution model. The implanted motifs and their JASPAR logos in each dataset are listed in Table S1.

### ChIP-seq datasets

We downloaded a total of 204 ChIP-seq datasets from Gene Expression Omnibus with accession number GSE61475(Tsankov et al. 2015), generated using a MNase-based ChIP-seq technique from early stages of endoderm (dEN), mesoderm (dME), ectoderm (dEC) and mesendoderm (dMS) tissues derived from human ES cells. Since motifs of some target TFs are not documented in the JASPAR database, therefore we excluded 99 datasets for these TFs from our analysis, resulting in a total of 105 datasets for 21 TFs with documented motifs in JASPAR (Table 4). We generated three groups of datasets G_1_, G_2_ and G_3_ by extracting 200, 500 and 1,000 bp, respectively, genomic sequence for each called binding peak in each dataset, with the summit of the binding peak being the center. We masked the repeat regions using Repeat Masker(Bedell et al. 2000) and Tandem Repeat Finder(Benson 1999) with the default parameter settings.

### Evaluation of the Programs

For each of predicted motifs, we compared it with motifs in JASPAR(Portalescasamar et al. 2010) using TOMTOM with Euclidean distance being the metric(Gupta et al. 2007b), and considered the best hit as a match if the *q*-value ≤0.05. To quantify the performance of the programs for identifying motif lengths, we computed three metrics: Performance Coefficient (PC), Positive Predictive Value (PPV) and Sensitivity (SN)(M et al. 2005; Ikebata and Yoshida 2015), based on the overlap between the predicted motif and the hit, defined as follows,

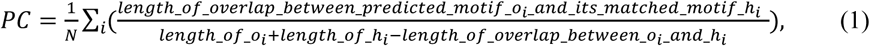

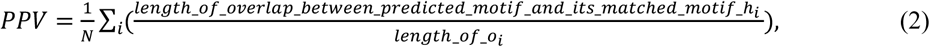

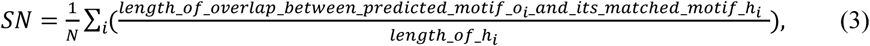

where *N* is the number of predicted motifs with a hit in JASPAR.

We present the results as mean ± standard error when appropriate, and compared the result of ProSampler with those of other programs evaluated using Wilcoxon rank sum test or two-tailed t-test as indicated in the text.

### The ProSampler Algorithm

#### Preprocessing

##### Step 1: Generating background sequences

For each ChIP-seq dataset, we generate a background sequence set with the same number and length of sequences using a third-order Markov chain model based on the frequencies of nucleotides in the dataset (Supplementary Methods) (Redhead and Bailey 2007).

##### Step 2: Identifying significant *k*-mers

We count all possible *k*-mers (*k*=8 by default) in both the ChIP-seq and background sequence sets, and at the same time record the two flanking *l*-mers (*l* =6 bp by default) of each *k*-mer. Let *n*_*F*_(*k*_*i*_) and *n*_*B*_(*k*_*i*_) be the counts of the occurrences of a *k*-mer *k*_*i*_in the ChIP-seq and background sequence sets, respectively. If search on both strands is desired, we combine the counts of each pair of reverse complementary *k*-mers. We evaluate each *k*-mer *k*_*i*_ for its significance using a two-proportion z-test with the null hypothesis that its frequencies in the ChIP-seq (*p*_*F*_(*k*_*i*_) and background sequence (*p*_*B*_(*k*_*i*_) sets are the same:

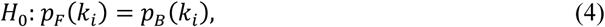

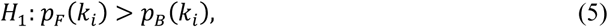

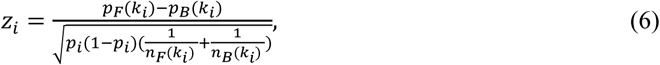

where,

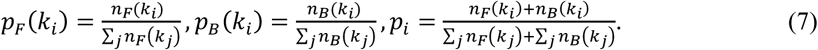

We consider that a *k*-mer *k*_*i*_ is significant or sub-significant if *z*_*i*_ is greater than a preset value *α* or *β* (*α* > *β*), respectively (by default, *α* = 8.0 corresponding to a *p*-value of 6.7 × 10^−16^, and *β* =4.5, corresponding to a *P*-value of 3.4 × 10^−6^). Let all the significant and sub-significant *k*-mers be the sets *K*_1_ and *K*_2_, respectively. Note that *K*_1_ is a subset of *K*_2_.

##### Step 3: Constructing preliminary motifs and their position weigh matrixes (PWMs)

For each significant *k*-mer *k*_*i*_∈ *k*_1_ we combine it with all other sub-significant *k*-mers 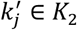 if their Hamming Distance(Forney 1992) 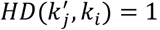, to form a preliminary motif. We construct its PWM *m*_*i*_ using the counts of the combined *k*-mers. Let *M* be the set of these PWMs. Notably, a *k*-mer can be included in multiple preliminary motifs. Then we sort the preliminary motifs according to their *z*-scores in the descending order.

##### Step 4: Constructing the motif similarity graph

We construct a graph using the PWM set *M* as the nodes, and connecting two nodes *m*_*i*_ and *m*_*j*_ if their Sandelin-Wasserman (SW) similarity(Sandelin and Wasserman 2004a; Gupta et al. 2007b) is greater than a preset value *γ* (by default, *γ* = 1.80), which is defined as

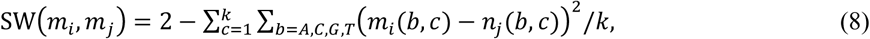

where *m*_*i*_ (*b,c*) and *m*_*j*_(*b,c*) are the frequencies of base *b* in column *c* of motifs *m*_*i*_and *m*_*j*_, respectively.

### Gibbs sampling

Our Gibbs sampler starts by combining the currently most highly ranked *m*_*i*_ (k_*i*_*’* s *z*-score) with its neighbors in the graph to form a seed motif *C*_*i*_ (**Fig. M1**) with redundant *k*-mers removed. In each cycle of sampling, we randomly select a motif *m*_*t*_ in *C*_*i*_, and then identify the motif *m*_*t*_ from the neighbors of *m*_*k*_that are not in 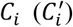 (**Fig. M1**) with the highest SW similarity to *C*_*i*_with *m*_*h*_removed. We add *m*_*t*_to C_*i*_, if the resulting *MotifScore(C*_*i*_+*m*_*t*_) is the better than *MotifScore(C*_*i*_-*m*_*k*_+*m*_*t*_)(replacing *m*_*h*_by *m*_*t*_) and *MotifScore(C*_*i*_)(the original score); we replace *m*_*h*_ by *m*_*t*_ in *C*_*i*_ if the resulting *MotifScore(C*_*i*_*m*_*k*_+*m*_*t*_) is the better than *MotifScore(C*_*i*_+*m*_*t*_)and *MotifScore(C*_*i*_) (the original score);we replace *m*_*k*_ by *m*_*t*_ in *C*_*i*_ if the resulting *Moltifscore*(*C*_*i*_ - *m*_*k*_ + *m*_*t*_)is better than where *Moltifscore* (*C*_*i*_ + *m*_*t*_) and *Moltifscore* (*C*_*i*_),where

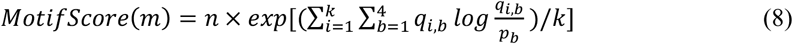

Where, *n* is the total count of the combined *k*-mers in motif *m,q*_*i,b*_the probability of base b appearing at position *i*, and *p*_*b*_the probability of base b in the dataset. We removed redundant *k*-mers in *C*_*i*_ after each updating.

After *N* cycles of iteration, we predict *C*_*i*_as a core motif with a length of *k*. We remove all the nodes in *C*_*i*_and inscribed edges from the graph to create a new smaller graph. We identify the next core motifs by repeatedly applying this process to the updated graphs until the graph becomes empty or a specified number of motifs are found.

**Fig. 7.**
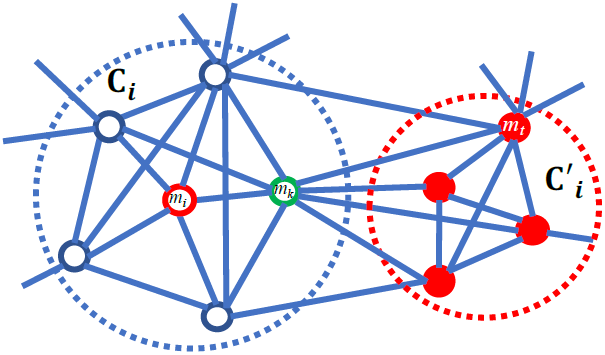
Schematic of Gibb’s sampling on the motif similarity graph. The most highly ranked preliminary motif *m*_*i*_and its neighbors form a seed motif *C*_*i*_ (open white nodes). In this cycle of sampling, a neighbor *m*_*k*_ of *m*_*i*_is randomly selected. All the neighbors of *m*_*k*_not in *C*_*i*_ (red solid nodes) are considered, and *m*_*t*_ is identified to have the highest SW similarity to *C*_*i*_ with *m*_*k*_ removed.

### Extending core motifs

To identify a motif longer than *k*, for each *k*-mer in each core motif, we pad its two flanking *l*-mers in the genome to the corresponding ends, extending the alignment with a length of 2 · *l+k*. We compare the frequencies of each nucleotide in each flanking columns starting from the closest ones to the core motif, with that in the dataset using the two-proportion z-test. We pad a flanking column to the core motifs if at least one of the column’s nucleotides has a significantly different frequency from that in the dataset (by default, z>1.96, corresponding to P-value < 0.05). We stop the extension in a direction once an insignificant column is encountered.

### Code availability

The source code, executives and relevant data are available under the GNU General Public License v3.0 at Github: https://github.com/zhengchangsulab/prosampler.

## ACKNOWLEDGEMENT

The authors would like to thank Dr. Meng Niu for assisting in preprocessing the original MNChIP-seq datasets, and members in the Su Lab for discussion, and members in the Li Lab for advices on manuscript writing. The work was partially supported by US National Science Foundation (DBI-1661332) and NIH (R01GM106013) to ZS, National Natural Science Foundation of China (61572358) and the Natural Science Foundation of Tianjin Science and Technology Committee (16JCYBJC23600) to SZ, and National Science Foundation of China (61432010, 31571354 and 61771009) to GL.

## AUTHOR CONTRIBUTIONS

Zhengchang Su conceived the original algorithmic idea and designed the algorithm. Dr. Guojun Li modified the algorithmic details of the algorithm subsequently. Yang Li and Pengyu Ni developed the software. In addition, Yang Li carried out all the computational experiments and analysis. Yang Li, Zhengchang Su and Guojun Li wrote the manuscript. All authors read and approved the final manuscript.

## COMPETETINGNG FINANCIAL INTERESTS

The authors declare no competing financial interests.

